# Alpha fluctuations regulate the accrual of visual information to awareness

**DOI:** 10.1101/2020.09.16.298166

**Authors:** Mireia Torralba, Alice Drew, Alba Sabaté San José, Luis Morís Fernández, Salvador Soto-Faraco

**Author notes:** <u>Corresponding author:</u> Mireia Torralba Cuello, ZIP: 08002.

## Abstract

Endogenous brain processes play a paramount role in shaping up perceptual phenomenology. This is illustrated by the alternations experienced by humans (and other animals) when watching perceptually ambiguous, static images. We hypothesised that endogenous alpha fluctuations in the visual cortex pace the accumulation of sensory information leading to perceptual outcomes. Here, we addressed this hypothesis using binocular rivalry combined with visual entrainment and electroencephalography in humans (64 female, 53 male). The results revealed a correlation between the individual frequency of alpha oscillations in the occipital cortex and perceptual alternation rates experienced during binocular rivalry. In subsequent experiments we show that regulating endogenous brain activity via rhythmic entrainment produced corresponding changes in perceptual alternation rate. These changes were observed only in the alpha range but not at lower entrainment frequencies, and were much reduced when using arrhythmic stimulation. Additionally, entraining at frequencies above the alpha range did not result in speeding up perceptual alternation rates. Overall, these findings support the notion that visual information is accumulated via alpha cycles to promote the emergence of conscious perceptual representations. We suggest that models of binocular rivalry incorporating posterior alpha as a pacemaker can provide an important advance in the comprehension of the dynamics of visual awareness.

## Introduction

There is ample agreement that our perception of the world is not built from incoming sensory input alone, but is also determined by endogenous brain processes (Friston, 2012). The impact of these endogenous processes is well illustrated by spontaneous fluctuations in conscious perception when watching ambiguous images, a phenomenon known as bistable perception (Sterzer et al., 2009). Ambiguous images lead observers to experience alternations between different perceptual interpretations of an otherwise static input and have been used by neuroscientists to investigate the neural correlates of fluctuations in perceptual awareness. Here, we address the hypothesis that alpha oscillatory activity in the occipital cortex plays a causal role in these perceptual fluctuations.

We focused on binocular rivalry (BR), a phenomenon that arises when two dissimilar images are presented to different eyes. BR has been characterised as an intrinsically dynamic process resulting from competition and mutual inhibition between different neural populations representing the rival images (Mamassian & Goutcher, 2005). There is still controversy about the specific brain areas engaged in this competition (Block, 2019; Leopold & Logothetis, 1999; Phillips & Morales, 2020), however, evidence suggests that the occipital cortex is involved in resolving perceptual ambiguity (De Jong et al., 2016; Lee et al., 2007; Polonsky et al., 2000). In some models, BR competition is not solely based on mutual inhibition but also on attentional modulation (Li et al., 2017).

Phenomenologically, BR is characterised by the natural alternation rate (NAR) between dominance periods, a stochastic process strongly dependant on properties of the rival stimuli (Jan W Brascamp et al., 2005; Kang, 2009; Sobel & Blake, 2002). Interestingly, the NAR is relatively stable within individuals and genetically hereditary (Miller et al., 2010). Additionally, although inter-individual variability in BR dynamics is large (Kleinschmidt et al., 2012), alternation rates within an individual correlate across different forms of bistable perception (Schmack et al., 2013) and stimuli (Carter & Pettigrew, 2003).

Recently, several studies have linked BR with brain oscillatory activity in different frequency bands (Doesburg et al., 2005; Piantoni et al., 2010), and in particular to occipital alpha activity (8-14 Hz) (Davidson et al., 2018; Katyal et al., 2019). Alpha oscillations are strongly related to attention (Klimesch, 2012; Thut et al., 2006) and, more relevantly, alpha activity in the occipital cortex is known to be related to rhythmic fluctuations in visual perception (Vanrullen et al., 2011). For example, the gating by inhibition hypothesis (Jensen & Mazaheri, 2010) contends that the alpha rhythm imposes a discrete sampling of sensory input via an alternation between moments of heightened neural excitability and bouts of pulsed inhibition, cycling every 100 ms approximately. In line with this hypothesis, variations in the speed of alpha oscillations correlate with inter-individual variability in the temporal resolution of visual perception (Samaha & Postle, 2015) and processing speed in general (Clark et al., 2004). The results of entrainment studies support the causal role of alpha oscillations in modulating perception (Cecere et al., 2015; Ronconi et al., 2018). An interesting property of the cortical alpha rhythm is that the individual alpha frequency (IAF) reflects systemic properties of the brain, is highly heritable, and relates to cognitive functioning (Grandy et al., 2013).

In this study, we focus on establishing a connection between the speed of alpha endogenous activity and typical BR alternation times. We propose that cortical alpha oscillations play a key role during the competition between alternative representations in BR. More specifically, we hypothesise that the alpha rhythm provides phasic windows for the cyclical re-evaluation of the competing stimuli representations and in consequence, evidence accumulation takes place in a phasic manner. This discrete, alpha gated, evidence accumulation eventually leads to perceptual switches. According to this hypothesis, faster alpha oscillations will correlate positively with the NAR in a BR task and visual entrainment will causally modulate BR dynamics in a frequency-specific way.

Up to date, the alpha gating by inhibition hypothesis (Jensen & Mazaheri, 2010) had not been considered in an account of perceptual alternations in Binocular Rivalry. The experiments presented here address, for the first time (to the best of our knowledge), the causal connection between perceptual alternations and alpha activity. We offer a mechanistic account of how visual information is accumulated, via discrete sampling in alpha cycles, to promote the emergence of conscious perceptual representations. The outcomes of this study help answering an outstanding question in perceptual awareness: What is the origin of individual variability regarding the speed of alternations in BR? If, as we put forward here, the origin of this variability can be traced down to alpha endogenous activity, models of BR incorporating alpha as a pacemaker can provide an important advance in the comprehension of the dynamics of visual awareness.

## Methods

### Subjects

Participants received 10€/ hour in return for their participation. They all provided written informed consent prior to the study and were naive to the purpose of the experiment. The study was run in accordance with the Declaration of Helsinki and the experimental protocol approved by the local ethics committee CIEC Parc de Salut Mar (Universitat Pompeu Fabra, Barcelona, Spain).

Subject inclusion criteria in all the experiments were: normal or corrected to normal vision, not being under medication and, for experiments including an electrophysiological recording (Experiments 1, 2 and 3), displaying a single peak in the alpha range during the eyes-closed pre-screening EEG recording (see *IAF estimation*). In order to minimise alternations unrelated to endogenous activity (Kanai et al., 2005), we excluded participants with a high percentage of EEG artefacts such as blinks, eye movements or motor contamination in general (with a high percentage defined as more than 5% in Experiment 1 and 10% in Experiments 2 and 3, applied separately in each of the 4 experimental conditions). For Experiments 2, 3, and 4 due to the reduced availability of data points per condition (5 blocks per condition compared to 9 in Experiment 1) we established an additional criterion to ensure a reliable estimation of the NAR (Supplementary materials and methods: *Inclusion criteria*): In Experiments 2, 3 and 4 we included only subjects with 50 or more percepts in each experimental condition which, according to simulation, ensured a deviation in median estimation <3.5%. Please note that in Experiment 1, all subjects had far more than 50 percepts (a percept is the period of time from seeing one pure colour to seeing the next pure colour).

Thirty-one participants (15 female, aged 18-34, mean age 23) participated in Experiment 1. Four participants were excluded for having more than 5% of the segmented epochs contaminated by visual and movement artefacts (final sample N=27). Twenty-nine participants (16 female, aged 18-31, mean age 24) participated in Experiment 2. One participant was excluded for presenting a double peak in the alpha range, two participants were discarded due to excessive artefacts and one participant was discarded due to insufficient time for estimation of the NAR (final sample N=25). Thirty-four participants (19 female, aged 19-32, mean age 24) participated in Experiment 3. One participant was excluded for presenting a double peak in the alpha range, two participants were discarded due to an insufficient number of percepts and one participant was discarded due to a technical problem with the recordings (final sample N=30). Forty participants (24 female, aged 18-34, mean age 24) participated in Experiment 4. Four participants were discarded due to an insufficient number of percepts and one participant was dircarded due to bad proportion of Green percepts (final sample N=35).

### Apparatus and Stimuli

Participants were presented with one Gabor patch to each eye, 11.5° in diameter, Michelson contrast 0.1, opposed orientations (±45° off vertical) and different colours (red, RGB= [1 0 0] and green, RGB adjusted individually to match the luminance of the red Gabor patch). Both patches were surrounded by a black annulus and contained a central fixation cross identical in each of the patches (Fig. 1 A). The images were presented on a 19.8-inch CRT monitor (120 Hz refresh rate) with a grey background (10.7 cd/m^2^) placed 80 cm from participants’ eyes. Each image was presented to a different eye by means of stereoscope mirror glasses (Fig 1 D). Visual stimuli were created and presented using the Matlab toolbox Psychtoolbox (RRID: SCR_002881), version 3.

**Figure 1:**
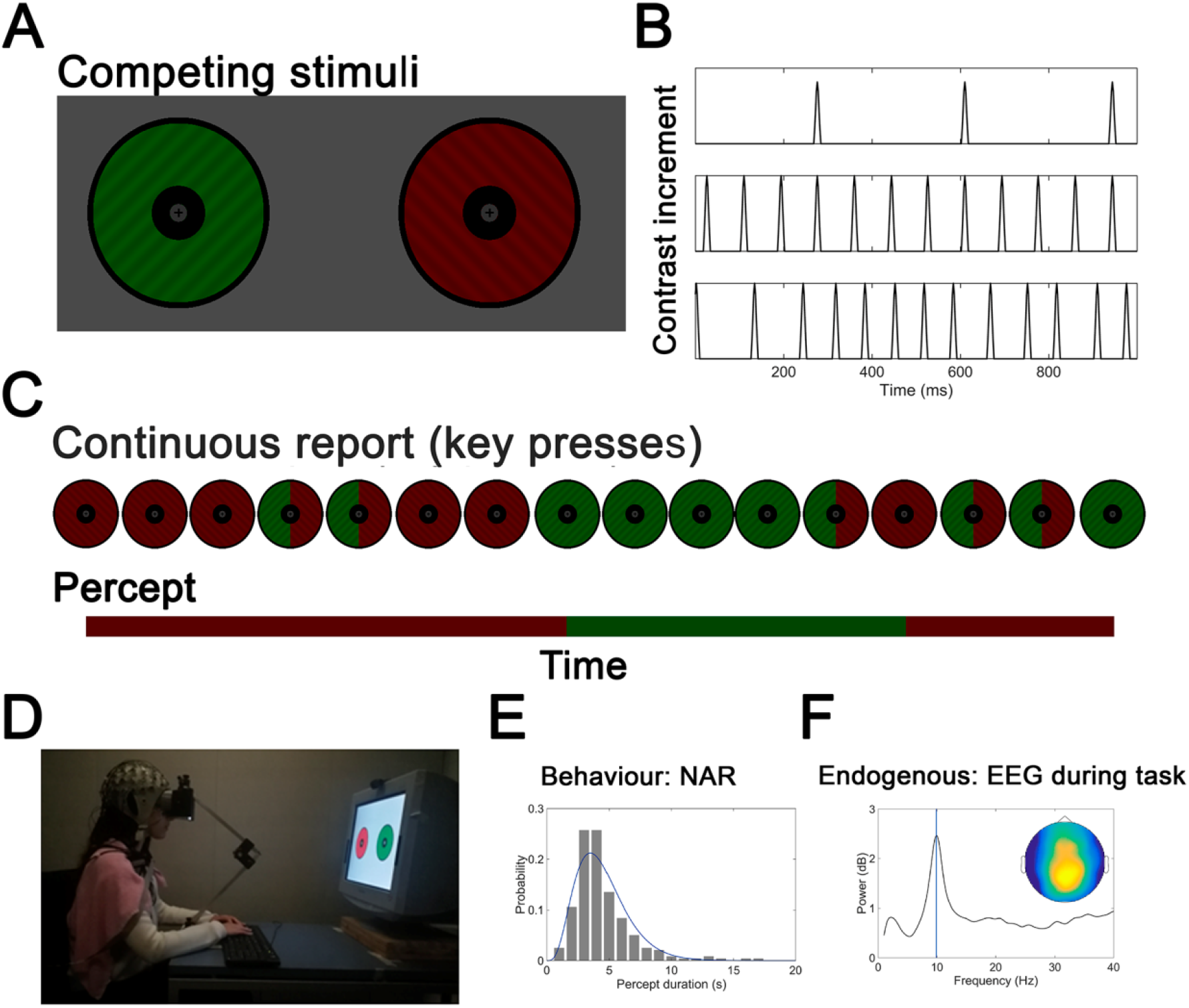
A: Example of the Gabor patches used in the experimental protocol. B: Contrast of both Gabor patches was modulated simultaneously with different temporal patterns (low and high frequency rhythmic modulation, fast arrhythmic modulation). C: Participants continuously reported the conscious percept (red, green, mixed or none) while EEG was concurrently recorded. D: Image of our experimental setup. E: Target behavioural measure was individual characteristic duration of conscious percept (NAR). F: Behavioural measures were complemented with endogenous oscillatory activity recorded during the BR task.

Visual stimulation for entrainment, when used, consisted of pulsed Gaussian increases in Michelson contrast from 0.1 to 0.6, of 58 ms duration in Experiments 2 and 3, and 41.16 ms in Experiment 4, presented simultaneously to the images displayed to the left and right eyes. Different entrainment frequencies were then achieved by varying the offset to onset time between pulses, whilst maintaining the slope of contrast increases constant (Fig. 1 B).

### Procedure

The procedure was identical for all experiments 1,2 and 3 (Supplementary Figure S1), with the only variation between experiments being the visual entrainment stimulation conditions (see below) and number of blocks performed. In experiment 4 we only collected behavioural data, and IAF at rest was not estimated, but the rest of experimental procedure was the same as in experiments 1, 2 and 3.

The experimental session started with the participant’s IAF estimation during a 5-minute rest block, with eyes closed. We then ran a protocol to match the two rival stimuli in terms of luminance for subjective equality (Cavanagh et al., 1987) (Supplementary materials and methods: *Objective isoluminance adjustment*). Next, the stereoscope mirrors were calibrated for the rival stimuli to appear at the same retinal location of each eye. Finally, two training blocks (120 s each) on the BR task were performed prior to the experimental BR runs.

All experiments contained the same rival images and were organised in 120 s blocks of a BR task. The eye of presentation of the red and green stimuli was alternated from block to block. After each block, there was a pause and participants were given the chance to rest before starting the next trial. The experimental conditions used in each individual experiment were characterised in terms of the visual stimulation used:

- No entrainment (0 Hz).
- Rhythmic entrainment: periodic contrast increments with a frequency of presentation of 3 Hz, 10 Hz, 17.14 Hz, IAF, IAF-2Hz and IAF+2HZ, depending on the condition.
- Arrhythmic visual stimulation (arrIAF): non-periodic contrast increments. In this condition, the latencies between contrast increments were drawn from an exponential distribution centred at subjects’ 1/IAF in order to match the number of contrast increments during the corresponding rhythmic IAF condition of the same subject.

In Experiment 1, subjects ran 9 blocks of the 0 Hz condition. In experiments 2, 3 and 4 subjects ran 5 blocks of each experimental condition: 3 Hz, IAF-2Hz, IAF and IAF+2Hz for Experiment 2 and 0 Hz, 3 Hz, IAF and arrIAF for Experiment 3. (Supplementary Figure S1)., and 0 Hz, 3 Hz, 10 Hz and 17.14 Hz for Experiment 4. The order of conditions was intermixed pseudo-randomly between blocks (Supplementary Figure S1).

During the task, participants were instructed to fixate their gaze on the central fixation cross and report their current percept (red, green, mixed or null) by key press with the left and right index of each hand (Fig 1 C). Response mapping for the green stimulus and the red stimulus was counter-balanced between participants. To report a mixed (piecemeal) percept, participants were instructed to press both keys simultaneously and to release both keys if none of the three aforementioned percepts were apparent (e.g., null percept).

### Predictions

In this study, we performed a series of four experiments, each of them designed to test specific predictions derived from our hypothesis. The predictions and associated experiments are as follows.

### NAR correlates positively with IAF

If alpha endogenous activity paces information accrual, one first expected result is that faster individual alpha oscillations will lead the individual to experience a faster natural alternation rate (NAR) in a BR task (Fig. 2). Experiment 1 was designed to test this prediction: EEG endogenous activity was concurrently recorded while participants performed a BR task. The experiment was pre-registered (https://osf.io/n5c8g/) before analysing any data. Recently, this prediction has been independently supported in a study showing a correlation between NAR and IAF (Katyal et al., 2019).

**Figure 2:**
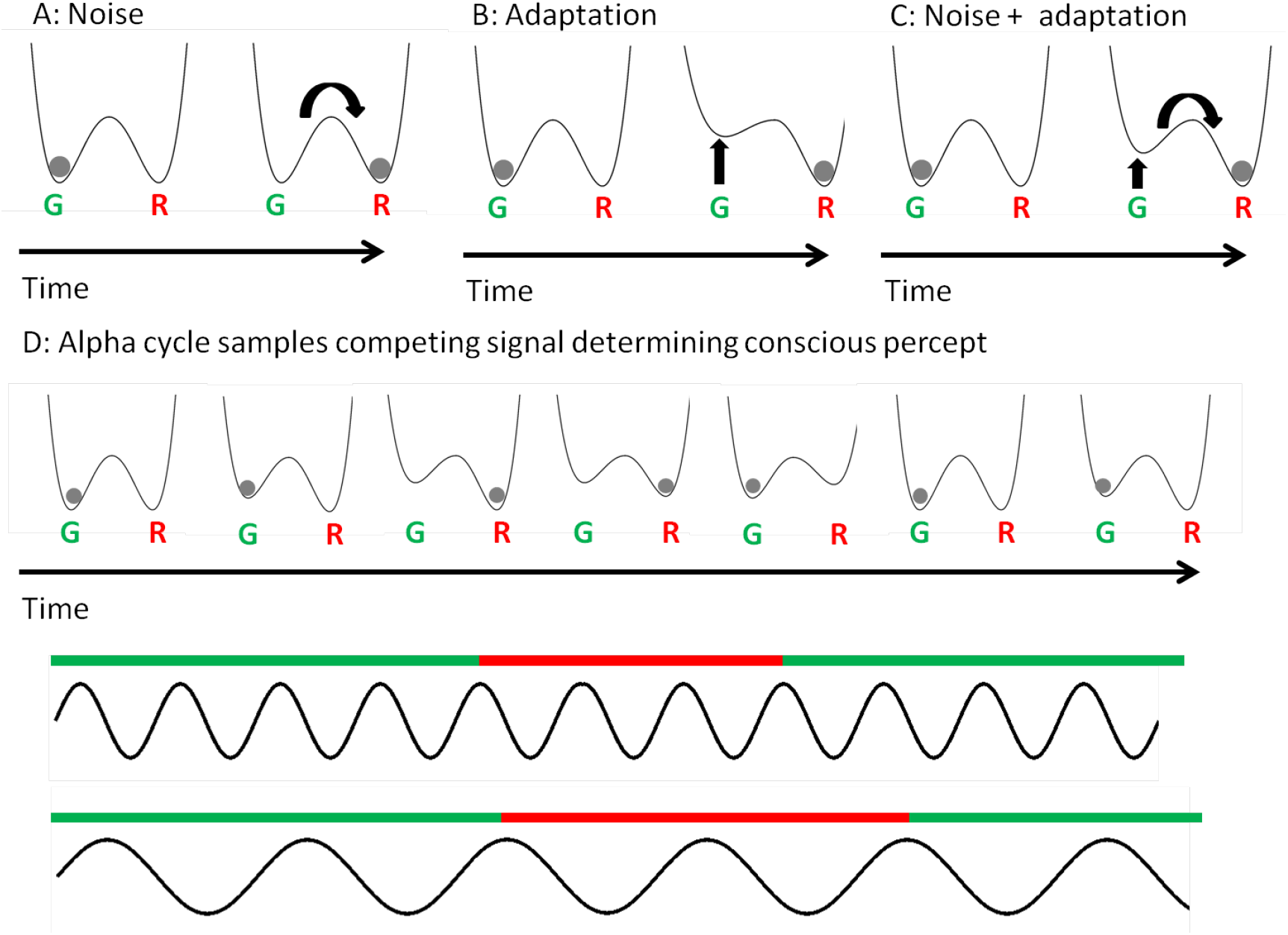
Binocular rivalry can be represented as a double well potential, each of the wells (here named R and G, for Red and Green respectively) represent the excitability of the competing neural populations (the larger the excitability, the deepest the well). The grey disk represents the neural population with stronger evidence. A: In some models, neural noise triggers perceptual switches, therefore the percepts could happen at any point in time. B: Other models predict that the excitability of the neural population sensitive to current conscious percept decreases over time, due to adaptation. As a result, as time decreases the probability of a switch increases, but the percept durations predicted by this model are highly irregular. C: At present, it is acknowledged that both noise and adaptation play a role in BR, therefore predicting stochastic percept durations (due to noise) and low probability of very long percept durations (due to adaptation). D: Top, sketch of a possible time course of the competition between neural populations during a BR task, as time evolves both noise and adaptation affect the instantaneous competition process. Middle: Alpha activity periodically samples the outcome of the competition process, and, as result, the conscious percept (colored line) is determined over time. Bottom: A slower alpha rhythm samples competition process more slowly and, as a result, perceptual switches are more spaced in time.

### Alpha activity is causal to BR dynamics

Furthermore, our hypothesis implies that alpha oscillatory activity is causal to BR dynamics. Therefore, we anticipated that entraining relevant endogenous alpha activity with rhythmic visual stimulation should cause changes in perceptual alternation rate. Based on this, our second prediction is that entraining brain activity slightly below the individual’s alpha rhythm will result in the individual experiencing slower BR alternation rates, whereas entraining brain activity slightly above the alpha rhythm will speed up alternation rates. Experiment 2 was designed to test this second prediction: we measured the NAR in a BR protocol while the observers’ brain activity was entrained at 3Hz, IAF-2Hz, IAF or IAF+2Hz.

### The effect of entrainment on BR dynamics is frequency specific and limited to periodic stimulation

One further implication of the hypothesis is frequency specificity. That is, only oscillatory frequencies neighbouring the alpha range are relevant to the re-evaluation of competing stimuli, and therefore entraining at rhythms outside the alpha range will not affect BR dynamics. Following this frequency specificity assumption, we predict that entraining brain activity at a frequency below the alpha range (e.g., 3 Hz) will not impact competition dynamics and, therefore, the alternation rates under non-alpha rhythmic entrainment will be equivalent to those without visual entrainment. For frequencies above the alpha band, the prediction is a bit different. Notice that, entraining at frequencies above the alpha range can result on the entrainment of a subharmonic frequency (Christoph S. Herrmann et al., 2016), therefore, it would be still possible to observe a change in the alternation rates in the fast (above alpha range) entrainment condition compared to the no entrainment condition. If a sub-harmonic is entrained, the alternation rates observed would be similar to those observed in the alpha range. However, we do not expect an increase in the alternation rate with respect to the alpha condition (e.g. 10 Hz). Therefore, we predict that entraining at frequencies above the alpha range will not result in faster alternation rates than those observed for entrainment at frequencies in the alpha range (e.g. 10 Hz).

We assume that that entraining with rhythmic Alpha activity (at or near IAF) will promote periodic sampling of the competing stimuli, by sustaining alpha activity over and above its naturally occurring ongoing dynamics, which is typically characterised by intermittent bouts (Rusiniak et al., 2018). According to this assumption, we expect that the rate of alternations in the alpha range will be faster when entraining near Alpha than in when entraining well below Alpha or in the no entrainment condition.

Finally, according to our hypothesis, the periodicity of neural fluctuations is consequential for BR dynamics. That is, modulations of BR dynamics imposed by entrainment depend on the temporal (rhythmic) structure of the stimulation and cannot be explained solely by accumulated changes in the amount of energy (i.e., increases in stimuli strength). Hence, our fourth main prediction is that the NAR in BR, observed under arrhythmic stimulation, will be slower than the NAR observed under rhythmic (alpha) entrainment. Experiment 3 was designed to test frequency specificity for low frequencies and periodicity. To test frequency specificity we compared the NAR while observers were entrained with 3Hz, with the one obtained under no entrainment (0 Hz). To test for periodicity we compared the NAR under rhythmic (IAF) entrainment to comparable arrhythmic stimulation. This experiment was pre-registered prior to any data analysis (https://osf.io/w8m4q/). Experiment 4 was designed to test frequency specificity further, including a test of low frequencies (0 Hz compared to 3 Hz) and high frequencies (10 Hz compared to 17 Hz). This experiment was pre-registered prior to any data analysis (https://osf.io/2yg85/).

For the sake of completeness, above we have formulated the final version of the predictions, revised after a reformulation of the hypothesis upon the outcome of Experiment 2 and before Experiments 3 and 4. This revision was included after we observed, in Experiment 2, an unexpected positive difference in NAR between IAF entrainment and 3 Hz entrainment that could, however, as we explain above, be still accounted for by our theoretical framework. After Experiment 2, we proceeded to a new formulation of the hypothesis (please refer to the registration form of Experiment 3 for more information; https://osf.io/w8m4q/), produced the adequate prediction, and designed Experiments 3 and 4 with specific tests of this prediction (that alpha entrainment would in fact promote the periodic re-evaluation of competing stimuli).

It should be emphasised that the aim of this study, outlined in these hypotheses, is merely to probe the role of alpha oscillations in rivalry dynamics. We make no claim as to the underlying cause of these fluctuations, which is a question beyond the ambitions of the present study. Regardless of their cause, perceptual fluctuations are a phenomenological effect of BR attributed to competition dynamics, and we set out to investigate the role of alpha oscillations in this visual competition. The mechanisms underlying perceptual fluctuations during BR have been addressed in a parallel study currently in preparation (Drew et al., 2019).

### Sample size estimation

The sample size for each of the three experiments was estimated to attain a 95% statistical power in the comparison of interest: IAF-NAR correlation in Experiment 1, IAF-2Hz compared to IAF+2Hz in Experiment 2, and IAF compared to arrhythmic stimulation in Experiment 3 (Supplementary materials and methods: *Sample Size estimation*).

### Behavioural analysis

Percepts shorter than 300 ms (estimated latency of the motor evoked potential from key presses (Halgren, 1990)) were filtered out from the analysis. The NAR was obtained as the inverse of the median time between reporting two different pure percepts (complete transitions from red to green or green to red) (see the distribution of percept durations for a representative subject in Fig. 1 E).

### EEG recording

EEG data were acquired using active electrodes (actiCAP, BrainProducts GmbH, Munich, Germany) with Brain Vision Recorder (BrainProducts GmbH, Munich, Germany) at a sampling rate of 500 Hz. The ground electrode was placed at AFz and online reference on the tip of the nose. Four external electrodes were used: left and right mastoid for offline re-referencing and horizontal and vertical electrooculogram for monitoring eye movements and blinks. Impedance was kept below 10 kΩ. In Experiment 1, 64 electrodes placed according to the 10-10 system were used for the recording. In Experiments 2 and 3, we recorded from a subset of 21 electrodes. The ROI (parieto-occipital electrodes), defined a priori, consisted of a cluster of 8 electrodes: PO9, PO3, POz, PO4, PO10, O1, Oz, O2 in Experiment 1, PO7, PO3, POz, PO4, PO8, O1, Oz, O2, in Experiments 2 and 3.

### EEG preprocessing and analyses

#### Pre-processing

EEG preprocessing and analysis was performed using Matlab’s toolbox Fieldtrip (RRID: SCR_004849) and custom-written code. The EEG data was band-pass filtered 1-70 Hz (Butterworth filter, order 2, two-pass); a notch filter was used to remove the 50 Hz line-noise. The data recorded for IAF calculation with eyes closed was processed without further analysis. The EEG data recorded during task performance was further visually inspected to mark visual or muscular artefacts. For the IAF measurement during the task (Experiments 1 and 3), data fragments containing artefacts were discarded from the analysis; for the evoked activity measurement (Experiments 2 and 3), it was not (please note that only participants with a very low percentage of artefacts (<10%) were included in the final sample). All analyses were performed in the parieto-occipital region of interest only.

#### IAF estimation

The IAF at rest was measured from continuous EEG recordings (300 s) using the Welch method (1 second segments, 10% overlap, 0.25 to 40 Hz in steps of 0.25 Hz). The power spectrum was averaged across the electrodes of interest (see ROI above) and background noise was estimated by fitting a 1/f^α^ curve to the power spectrum (Haegens et al., 2014). IAF at rest was used only as a screening: participants presenting a single peak in the spectrum exceeding background noise in the 5 to 15 Hz range were accepted to proceed with the experiment. The calculation of IAF during task was performed in the 0 Hz (no entrainment) condition (Experiments 1 and 3) (see the power spectrum of EEG activity during task for a representative subject in Fig. 1 F). Data were segmented into epochs according to participants’ reports (green or red percept, between key presses to avoid motor contamination). Pre-processed epochs were subsequently divided into non-overlapping segments of 1 s duration, demeaned and zero-padded to 10 s (Haegens et al., 2014). The power spectrum was calculated using a STFT (Hanning window, 4 to 30 Hz in steps of 0.1 Hz) and background noise correction using a 1/f^α^ fit was performed as in the estimation of IAF at rest (see above).

#### SSVEP and phase-locking measures

In order to determine that endogenous activity was indeed entrained during our experiments we performed several tests: we calculated the power spectrum of steady-state visual evoked potentials (SSVEP) for all the rhythmic conditions (Experiments 2 and 3) to test that we observed a peak of activity at the entrained frequencies, both at group and individual level (Supplementary materials and methods: *Entrainment at single subject level*) and also verified the degree of phase locking of neural signal to external stimulation.

SSVEPs were obtained for each of the rhythmic entrainment conditions by aligning the 5 blocks of each entrainment condition (four conditions in Experiment 2, and two in Experiment 3) to stimulus presentation. SSVEPs were then averaged across the electrodes of the ROI and the power spectrum was calculated with the same method described above for the IAF at rest, but using a window length of 2 seconds (in order to capture at least 5 cycles of the slowest entrainment activity, 3 Hz, in each of the windows of analysis). We normalised frequencies relative to the IAF prior to averaging the power spectrum across participants.

To calculate phase locking we used cross-coherence (XCOH) (Keitel et al., 2017) to assess the extent to which neural activity was synchronised to rhythmic stimulation. We calculated XCOH between external rhythmic stimulation (train of Gaussian contrast increments) and each of the channels of the ROI for each of the blocks (we discarded the initial and final 5 seconds of each block, resulting in 110 s of continuous data). We used 1 second non-overlapping segments (Hanning window) and 2048 points to calculate the FFT. Coherence was averaged across trials and electrodes of the ROI for each participant and experimental condition. Frequencies were normalised to IAF cycles and, finally, the value of coherence at each of the entraining frequencies of interest was selected and averaged across participants.

For the IAF condition in Experiment 3, we applied an additional entropy analysis adapted from Notbohm et al. (Notbohm et al., 2016), in order to show that rhythmic stimulation results in a stronger phase synchrony than arrhythmic stimulation. The time series of visual stimulation and concurrently recorded EEG were band-passed around the frequency of stimulation (see above) for rhythmic and arrhythmic conditions. Segments of data containing artefacts were discarded. For clean data, we obtained the phase difference between visual stimulation and concurrent EEG and calculated modified entropy as described by Notbohm et al. (Notbohm et al., 2016), using N=80 bins for entropy calculation. In case of entrainment, modified entropy would be larger for the rhythmic condition compared to the arrhythmic condition.

### Pre-registered statistical tests

For all statistical tests, the alpha level was set to 0.05. The correlation between the NAR and IAF was tested by means of a Pearson correlation (right tailed). The impact of alpha speed on the NAR was tested by means of a paired t-test (left tailed) for IAF-2Hz compared to IAF+2Hz. A one tailed t-test (right) was used to test the hypothesis that IAF entrainment results in a faster NAR than arrhythmic entrainment. A one tailed t-test (left) was used to test that entrainment at frequencies above alpha range (17.14 Hz) did not result in faster alternation rates that entrainment at frequencies in the alpha range (10 Hz).

A “two one sided tests” (TOST) (Lakens, 2017) was used to assess frequency specificity (following the prediction that only entrainment at or around alpha impacts BR dynamics). A TOST is significant when both p-values for left and right differences are below the set significance alpha level and the CI is contained within the pre-defined equivalence interval. For this test, the NAR at 3 Hz was normalised with respect to the 0 Hz NAR, and the equivalence interval was set to [-10 10] % before collecting data for Experiment 3 (https://osf.io/w8m4q/) based on simulations (Supplementary Materials and methods).

### Exploratory and control statistical tests (not pre-registered analyses)

In order to extend the evaluation of the potential correlation between IAF and NAR across different datasets, the p-values from correlations in Experiments 1 and 3 were combined by means of the Stouffer method (VanRullen, 2016).

The entrainment measure for Experiments 1 and 3 (EEG phase-alignment to external stimulation) was assessed by testing the significance of XCOH peaks at each of the entrainment frequencies. We applied the statistical test described by Biltoft et al (Biltoft & Pardyjak, 2009) using 2048 points for FFT estimation, 5 blocks of data, and 110 seconds of duration each at 500 Hz sampling rate, which resulted in 147 degrees of freedom. In this range of parameters, any coherence peak larger than 0.0405 would be significant.

Lastly, the comparison regarding neural signal synchrony between rhythmic and arrhythmic stimulation conditions in Experiment 3 was assessed by means of a paired one-tailed Wilcoxon test (Fay & Proschan, 2010).

## Results

According to behavioural responses in the BR task, the proportion of time spent in each of the two rival percepts was balanced in all three experiments. On average, participants reported a red percept 44±8% of the total recorded time, green 50±7% and no percept 6±6%, with similar proportions for all experimental conditions. For the calculation of the NAR, we used, on average, 231±71 percepts per condition for Experiment 1, 179±66 for Experiment 2 and 171±85 for Experiment 3.

Although IAF at rest can sometimes differ from IAF measured during task (Haegens et al., 2014), we did not observe these differences here for Experiment 1 (IAF_rest_ – IAF_task_=0.14±1.1 Hz, t(26)=0.77, p-value=0.4) nor for Experiment 3 (IAF_rest_-IAF_task_=0.14±0.9 Hz, t(29)=1.30,p-value=0.2). The differences were evaluated by means of a paired t-test (two-tailed). In Experiment 2, as all conditions included entrainment, IAF could not be measured during the task without interference and this test was not performed.

### The speed of endogenous alpha activity correlates with the speed of perceptual alternations

In line with our first prediction, in Experiment 1 we observed a significant correlation between IAF during the task and NAR (Fig. 3) (r=0.35, p-value=0.037). Visual inspection of the correlation plot opened the question of whether significance might be driven by two potential outliers with low IAF frequencies. From an electrophysiological point of view, the IAFs of these two subjects fall within the usual range of IAFs reported in the literature, 7 to 14 Hz (Haegens et al., 2011; Smulders et al., 2018), and therefore provide informative points in the analysis. Nevertheless, according to the Matlab toolbox Robust Correlation (Pernet et al., 2013), these two points were classified as bivariate outliers. In order to discard that the significant correlation observed in the main analysis was due to outliers, we calculated the robust correlation coefficient with the above-mentioned toolbox, following the recommendations of Pernet et al (Pernet et al., 2013). Data were distributed normally (Henze-Zirkler normality test p-value = 0.21) and included two bivariate outliers. We used the skipped Pearson correlation, which estimates the strength of the linear correlation ignoring outliers but taking into account the overall structure of the data, as recommended for this situation (Pernet et al., 2013). The correlation parameters obtained were in line with those obtained with the main (pre-registered) analysis (r=0.35, p-value=0.037).

**Figure 3:**
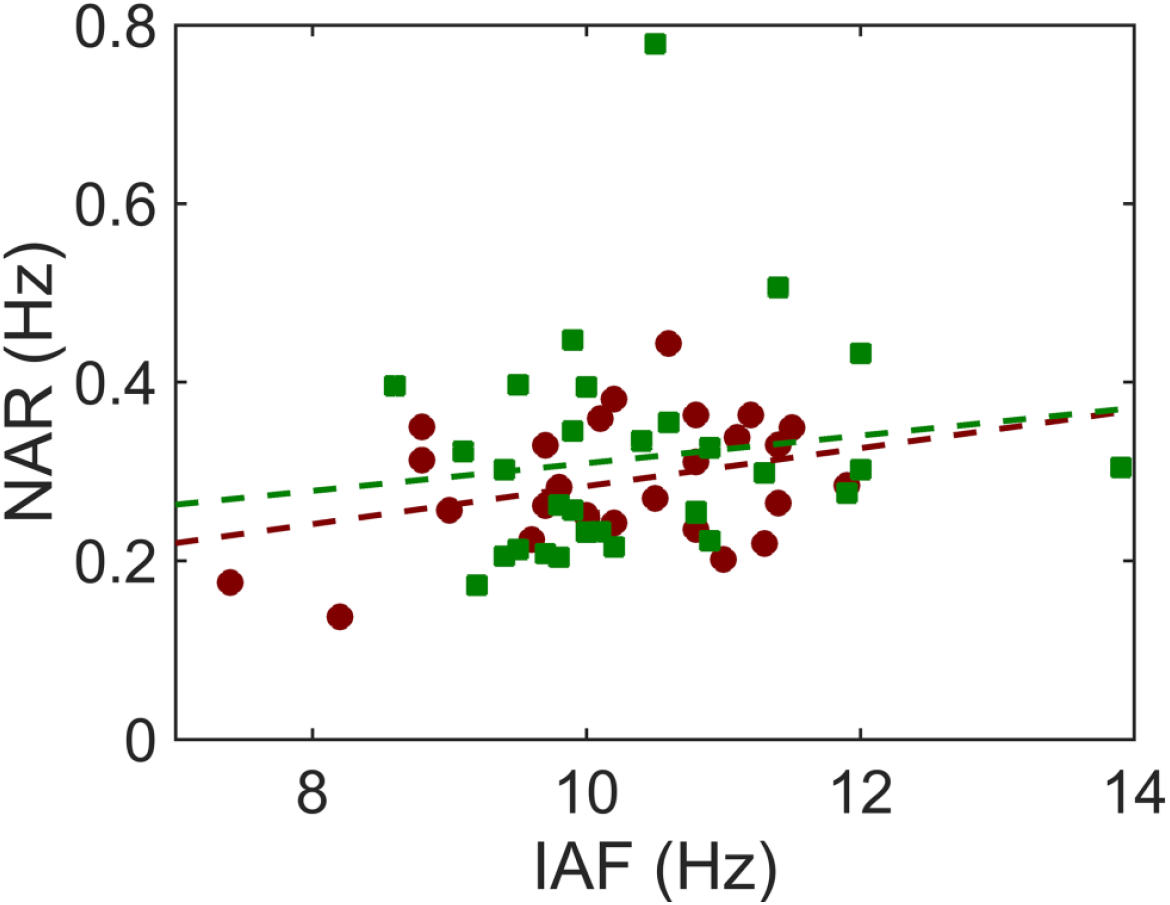
Natural alternation rate as a function of IAF during task for Experiments 1 (red) and 3 (green). Dotted lines correspond to the linear fit of NAR as a function of IAF.

Note that Experiment 3, which was designed to test the effect of periodicity and frequency specificity, also contained a no-stimulation condition, and therefore afforded an estimation of the correlation between IAF and NAR, but with substantially fewer trials. Here, we observed a correlation in the expected direction but that was not significant (r=0.19, p-value=0.15), perhaps not surprisingly given the reduced statistical power. As the dataset in Experiment 1, data in Experiment 3 were distributed normally (p-values for the Henze-Zirkler normality test p-value = 0.21) and included six bivariate outliers. The skipped Pearson correlation returned a similar estimate (r=0.23 p-value=0.11 for Experiment 3).

Taken individually, the evidence for the correlation between IAF and NAR supports the initial hypothesis, but it is not particularly strong. In order to garner more support, we used combined statistics and meta-analysis, in post-hoc confirmatory analyses. First, we found that the combined statistics of Experiment 1 and 3 (Stouffer method) rendered a significant effect (p-value=0.024). In addition, we performed a meta-analysis using the four independent datasets that, as far as we know, have tested IAF-NAR correlation: (Katyal et al., 2019), data from a previous experiment of our group (Pápai et al., 2018), and data from Experiments 1 and 3 in the current study. The meta-analysis was performed with the R package “meta”, using Fisher’s z transformation of correlations. In order to homogenise datasets between studies we transformed all behavioural data to natural alternation times (NAT=1/NAR), the method used by Katyal et al. (please note that contrary to the NAR-IAF correlation, the expected direction for the NAT-IAF correlation is negative). For the same reason, we used the mean of the percept duration distributions instead of the median. This meta-analysis provided further confirmation of the correlation between BR dynamics and endogenous IAF activity (r=-0.44, p-value<0.001). Lastly, in order to take into account publication bias in the meta-analysis, we used the p-uniform method (Studies, 2014). This method assesses the distribution of the sum of independent uniformly distributed random variables using the p-method (Irwin-Hall distribution, as implemented in the p-uniform package for R https://cran.r-project.org/web/packages/puniform/). After correcting for publication bias, the correlation was, once more, confirmed (r=-0.47, p-value=0.011). Altogether, the data supports our first hypothesis regarding the prediction that alpha frequency is correlated with the natural alternation between percepts, across individuals.

### The frequency of endogenous activity is causally related to the rate of alternations in conscious perception

In order to make sure that rhythmic visual stimulation successfully entrained endogenous oscillatory activity, in Experiment 2, we evaluated SSVEP and cross coherence (XCOH) at group level. The group-averaged SSVEP displayed a peak in the corresponding frequency of each of the entrainment conditions (Fig. 4. A), and the cross-coherence measure presented a significant peak (3 Hz: XCOH=0.1097, p-value=0.0002; IAF-2Hz: XCOH=0.135, p-value=0.00003; IAF: XCOH=0.0841, p-value=0.0017; IAF+2Hz: XCOH=0.1133, p-value=0.0002) (Fig. 5. B and C).

**Figure 4:**
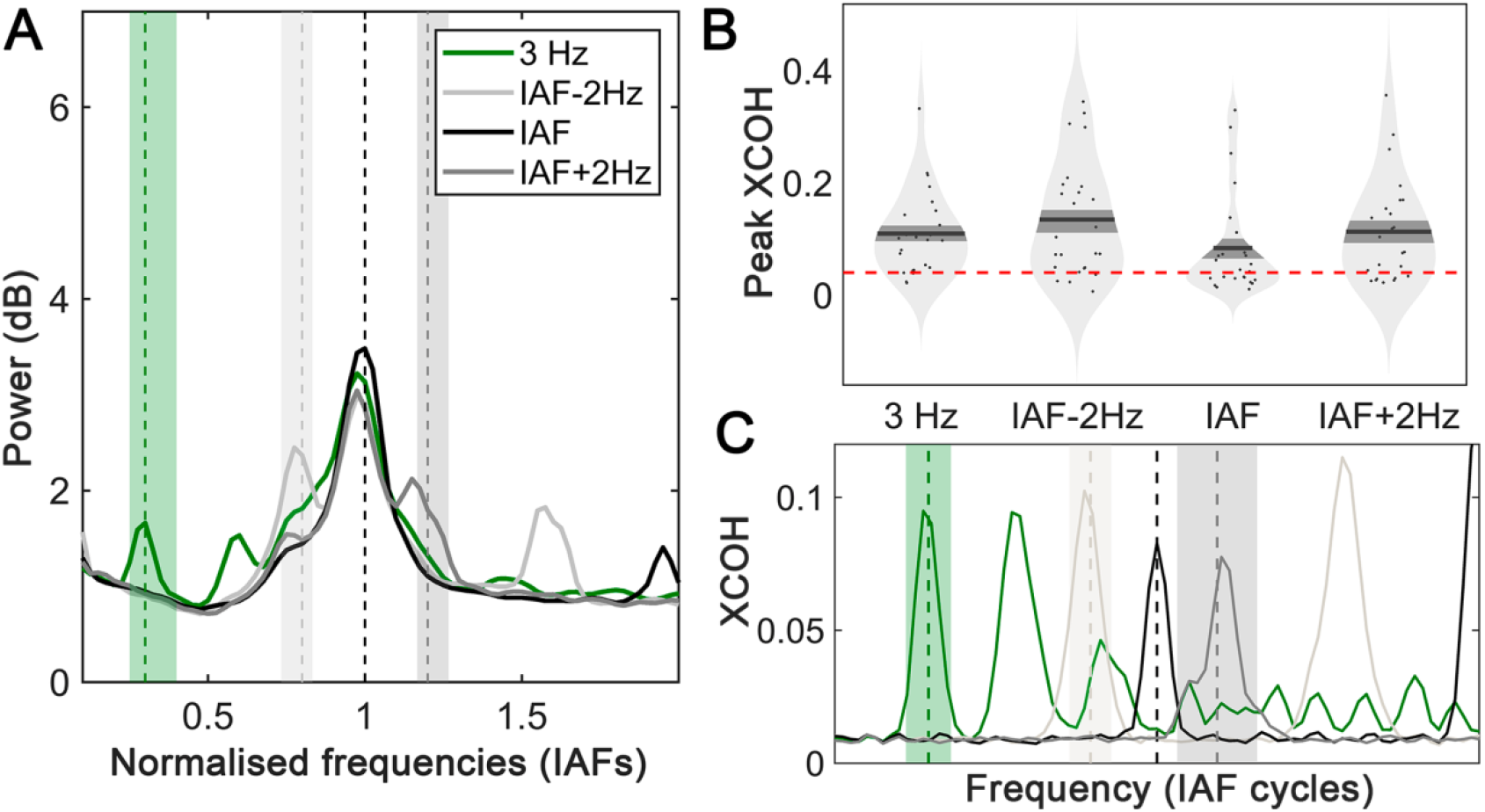
Entrainment in Experiment 2. A: Group average SSVEP. The frequencies are normalised with respect to rest IAF (the one used during IAF entrainment) and 1/f background noise correction was applied prior to the group averaging. Vertical dotted lines represent the target frequency at each condition (only IAF is exactly equal to one in the normalised plot, shaded areas indicate the lower and upper bounds for 3 Hz, IAF-2Hz and IAF+2Hz normalised values for each participant). B: Cross-coherence between the EEG signal and the rhythmic entrainment at the frequencies corresponding to each entraining condition. Dots represent individual values, black lines are the mean value for each condition and the dark shaded area corresponds to the standard error of the mean. The red dotted line presents the value of the coherence at a significance level 0.05. C: Cross-coherence spectrum for each of the experimental conditions.

**Figure 5:**
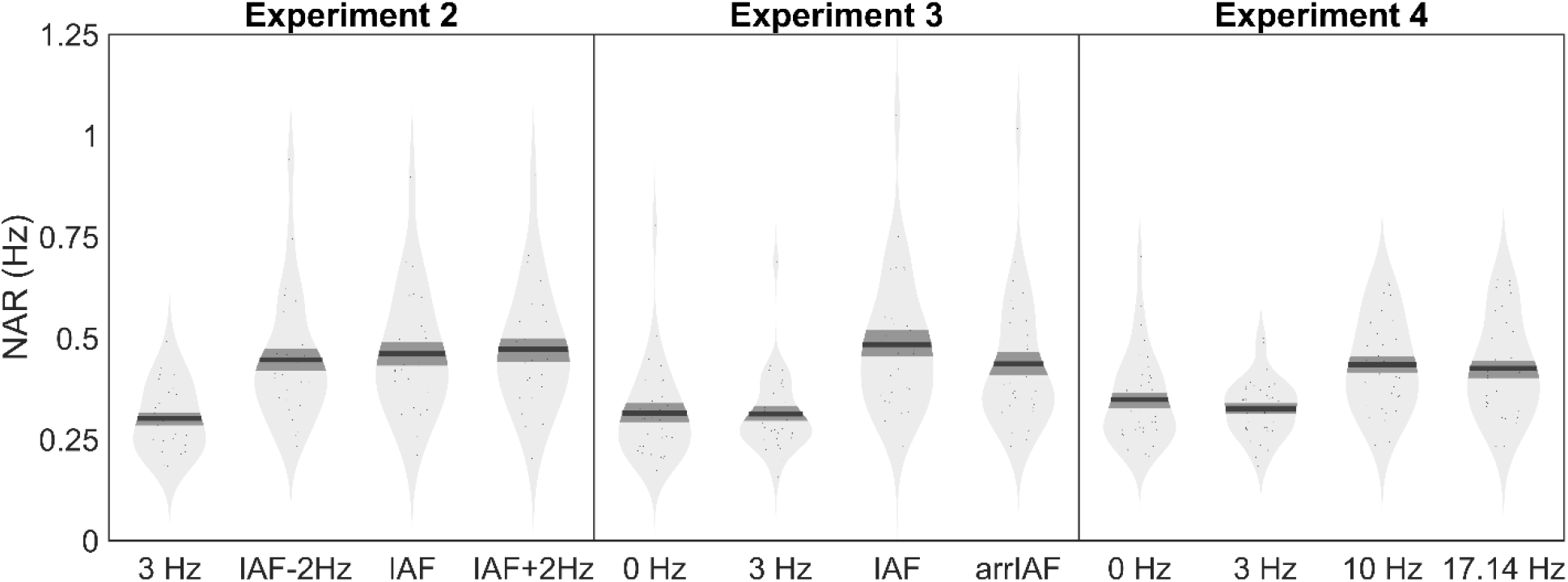
Effect of entrainment on perceptual alternations. NAR for Experiments 2, 3 and 4 at the different experimental conditions.

In Experiment 2, the NAR at 3 Hz was slower than any of the other conditions (Fig. 5. and Table 1). In the condition of interest, as expected, we observed a significant decrease in the rate of perceptual alternations during the slow alpha entrainment condition (IAF-2Hz) with respect to the fast alpha entrainment condition (IAF+2Hz) (p-value=0.03, t(24)=-1.942, d=0.39) (Fig. 6.). Entraining at the IAF produced NARs between the slow IAF-2Hz and the fast IAF+2Hz entrainment, as one would expect, but it was not significantly different from either of the two (Supplementary results: *Non-planned contrasts*). This finding confirms our second prediction, regarding the causal modulation of perceptual alternations by visual entrainment.

**Figure 6:**
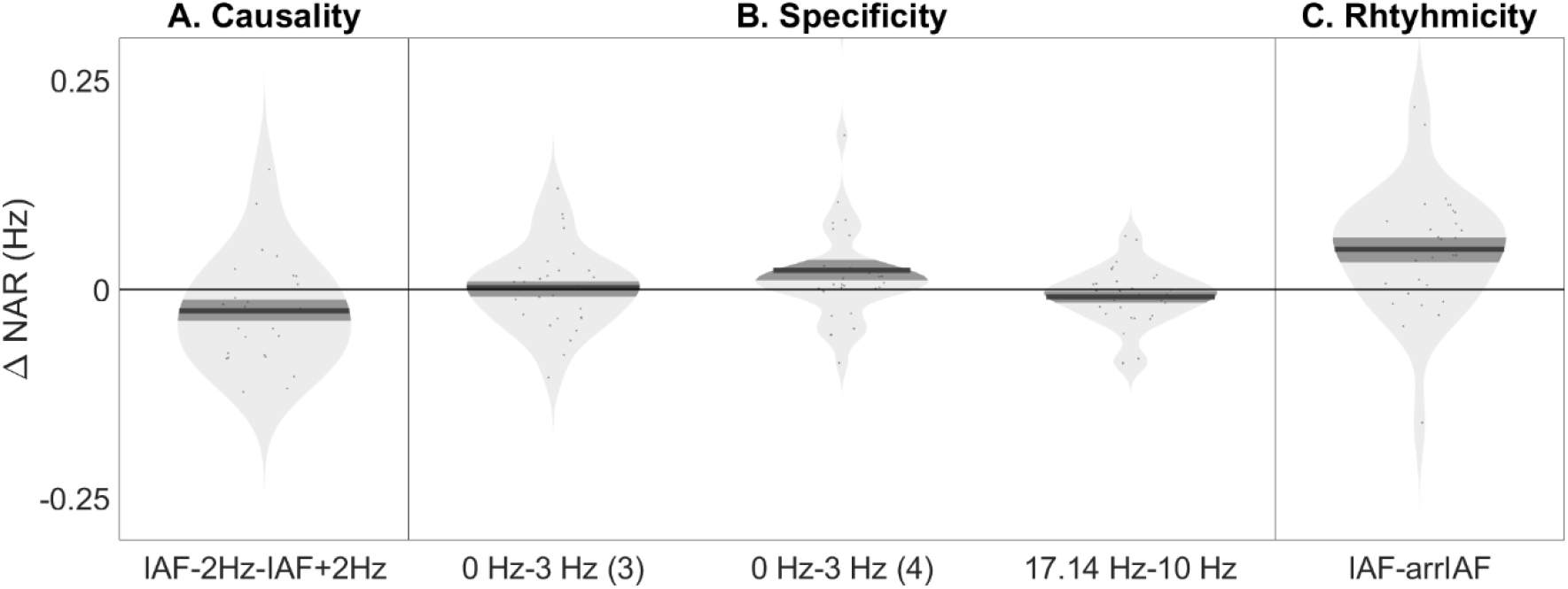
Comparisons of interest regarding each of the predictions. A: In experiment 2, a causal connection was observed between BR dynamics and endogenous activity. We observed faster alternations for fast alpha activity compared to slow activity (IAF-2 Hz compared to IAF+ 2Hz). B: In experiments 3 and 4, we verified that the effect was specific for alpha band. Alternation rates for entraining frequencies below the alpha range were equivalent to alternation rates without entrainment in both experiment 3 and 4 (3Hz vs 0 Hz). Additionally, in experiment 4, we observed that stimulating at frequencies above the alpha range did not result in alternations faster than the observed for alpha stimulation (17.14 Hz vs 10 Hz). C: In experiment 3, we verified that the speeding up observed in the IAF condition could not be fully explained by changes in the stimuli properties, but rhythmic stimulation resulted in faster alternation rates (IAF vs arrIAF)

### Rhythmic entrainment boosts the re-evaluation of competing stimuli

In Experiment 3, group averaged SSVEPs displayed peaks at each of the entraining conditions (Fig. 7. A). Endogenous activity was successfully entrained, as confirmed by the significant cross-coherence in all experimental conditions (3 Hz: XCOH=0.1251, p-value=0.0001; IAF: XCOH=0.0711, p-value=0.0048) (Fig. 7. B and C) and the higher values of normalised entropy observed in the IAF compared to arrIAF condition (p-value=0.000001) (Fig. 6. D). As would be expected, the cross-coherence for the arrhythmic condition was not significant (Fig. 6. B and C) (arrIAF: XCOH=0.016 p-value=0.32).

**Figure 7:**
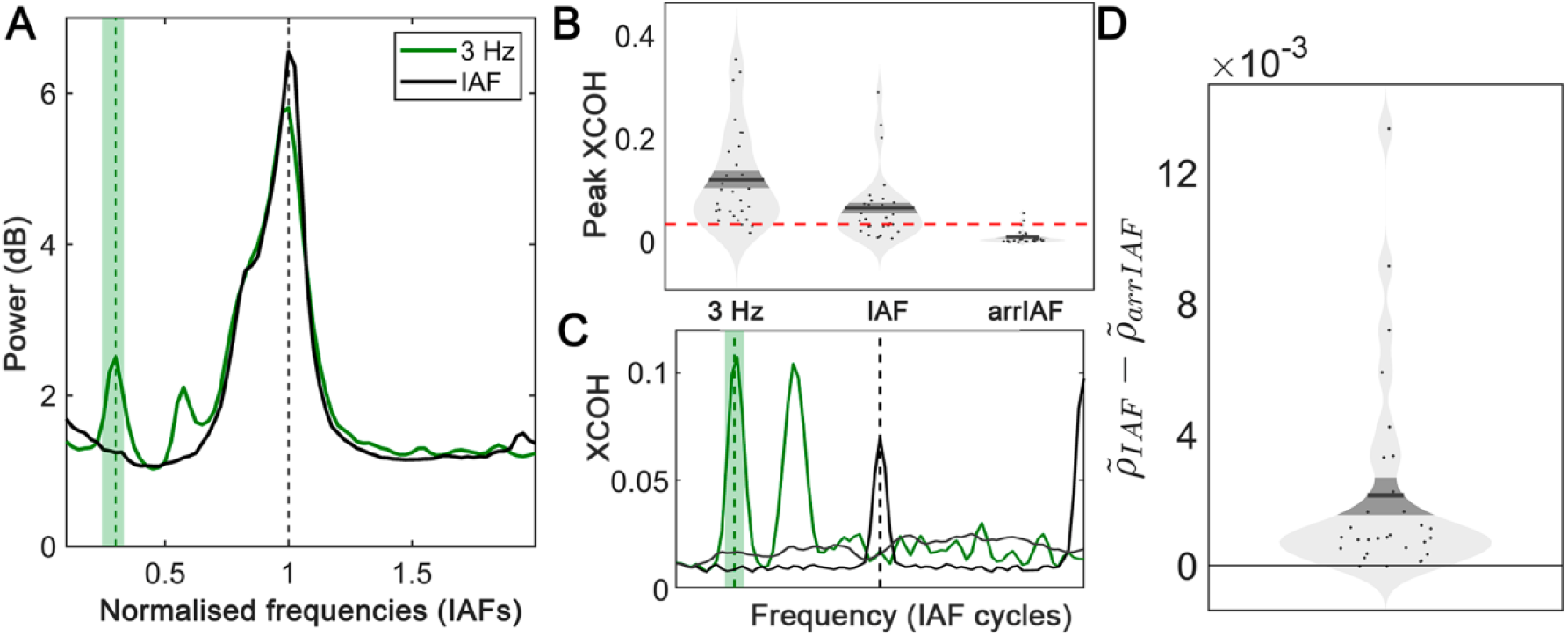
Entrainment in Experiment 3. A: Group average SSVEP. The frequencies are normalised with respect to rest IAF (the one used during IAF entrainment) and 1/f background noise correction was applied prior to the group averaging. Vertical dotted lines represent the target frequency at each condition (only IAF is exactly equal to one in the normalised plot, shaded area indicate the lower and upper bounds for 3 Hz, normalised values for each participant). B: Values of coherence between neural signal and rhythmic stimulation at the frequencies corresponding to each entraining condition. Dots represent individual values, black lines are the mean value for each condition and the dark shaded area corresponds to the standard error of the mean. The red dotted line presents the value of the cross-coherence at a significance level 0.05. C: Cross-coherence spectrum for each of the experimental conditions. D: Difference between normalised entropy at IAF and arrIAF conditions. Dots represent individual values, black thick line is the mean value and dark grey shaded area is the standard error of the mean.

The comparison of interest here was between the rate of perceptual alternations during rhythmic (at IAF) and arrhythmic (arrIAF) entrainment. In the arrhythmic condition, intervals between pulses were randomly picked from an exponential distribution of latencies centred at the inverse of IAF. IAF entrainment resulted in significantly faster NAR than arrhythmic entrainment (p-value=0.0005, t(29)=3.659, d=0.67) (Fig. 6), indicating that periodicity plays a key role in BR dynamics. This confirms the third main prediction, regarding periodicity.

In other words, although arrhythmic and rhythmic alpha entrainment always resulted in faster perceptual alternation rates compared to low-frequency (3 Hz) or no visual entrainment (0 Hz) (Fig. 6 and Supplementary results: *Non-planned contrasts*)), the increase in alternation rate in IAF entrainment cannot be solely attributed to an increase in the number or rate of pulses per unit of time (J W Brascamp et al., 2015). Instead, a mechanism involving periodicity is required to fully explain the result.

### Modulations in perceptual alternation rate are specific to entrainment in the alpha band

Further confirming the frequency-specificity implication of the hypothesis, we observed that the speeding up of perceptual alternations was constrained to the alpha band. Conditions that did not involve stimulation in the spectral vicinity of alpha always produced slower perceptual alternation rates. What is more, 3 Hz entrainment produced an alternation rate equivalent to no stimulation at all, as measured with two one sided tests (TOST (Lakens, 2017)) for an equivalence interval of 10% in both Experiment 3 (p-value_left_=0.007, p-value_right_=0.0001, CI= [-0.07 0.03]) (Fig. 6) and Experiment 4 (p-value_left_=2·10^−6^, p-value_right_=0.0071, CI= [-0.01 0.08]). Importantly, we did not observe a faster alternation rate when entraining at frequencies above the alpha range (p-value=0.9, t(34)=1.513, d=0.26)), thus confirming the frequency specificity.

## Discussion

We addressed the new hypothesis that alpha fluctuations in the occipital cortex mediate the accumulation of visual information, leading to the emergence of perceptual representations. Our findings, so far, seem to stand in good agreement with this hypothesis. First, we confirmed the direct prediction of our hypothesis that the speed of fluctuations in cortical alpha activity correlates with the dynamics of perceptual fluctuations in BR, a result also observed in another independent study (Katyal et al., 2019). The evidence provided by each individual experiment was weak to moderate, but aggregated results from four experiments indicate a robust relationship between Alpha frequency and alternation rate. The hypothesis further proposes that the endogenous alpha rhythm provides temporal windows for the re-evaluation of competing visual representations based on visual input. Consequently, we predicted that inducing faster endogenous activity should accelerate perceptual switches, a result we observed in Experiment 2. Moreover, our results suggest a causal relationship between alpha activity and awareness dynamics, because modulation of endogenous brain rhythms resulted in consequential changes in conscious perception. That is, speeding up or slowing down alpha cortical activity with entrainment alters the rate of perceptual alternations accordingly. Critically, this causal modulation is restricted to the alpha range and its effectiveness is subject to the periodicity of stimulation.

These findings are consistent with the pacemaker role of alpha activity in perceptual alternations, which we advance in the present hypothesis. The proposal builds on two well-known principles. First, that reciprocal inhibition between the neural populations selective of the competing stimuli is the mechanism causing perceptual alternations in BR (Tong et al., 2006). Second, that the activation of the neural population selective of the dominant conscious percept tends to decrease over time due to adaptation (Shpiro et al., 2009), making perceptual switches more likely with time. In this context, inhibition plays an active role: by stifling the neural population with lower excitability, it increases the signal to noise ratio corresponding to the alternative population (Klimesch, 2012). What we hypothesise is that this process takes place in a phasic manner: each alpha cycle opens a window for re-evaluation of the competing neural signals. In each of these windows, the weight of each neural population is evaluated, and the outcome of each evaluation determines the resulting conscious percept. Successive alpha cycles lead to the progressive accumulation of evidence toward one or the other percept.

Could this accumulation happen with changes in excitability regardless of rhythmic fluctuations? Levelt’s propositions predict that an increase in stimulus strength will result in faster perceptual alternations (J W Brascamp et al., 2015). In Experiment 3, we tested whether local, non-rhythmic changes in stimulus contrast could account for the increase in perceptual alternation rate during entrainment. Although arrhythmic stimulation increased alternation rates with respect to no-stimulation, these changes could not account for the effect in full. Rhythmic IAF stimulation resulted in faster alternations than arrhythmic stimulation. Additionally, if local changes in contrast had been responsible for speeding up switch dynamics, a difference between the no-stimulation condition and 3 Hz stimulation condition would have been expected. Yet, these two conditions rendered equivalent, and low, alternation rates.

Our manipulation could have affected the interplay between noise and adaptation, reflected in the coefficient of variation. Studies using anti-phase stimulation have reported drops in the coefficient of variation (Kim et al., 2006), implying a decrease in the contribution of noise to BR dynamics. In our study, the analysis of the coefficient of variation implied that the visual stimulation protocol used here did not modify the basic mechanism of neural competition. Therefore, one can assume that the competition mechanism remained unchanged and that only the frequency and probability of windows favouring the re-evaluation of perceptual representations were modified.

Sensory entrainment at or around the IAF produced a dramatic increase in perceptual alternations compared to 3Hz or no-stimulation. That is, alpha entrainment produces a sizable effect, even if there is already an ongoing rhythm at that frequency. This result was initially not expected, and motivated a reformulation of our initial hypothesis (please see Predictions section above) to account for the origin of this increase. In line with the well-known fact that endogenous alpha occurs in bursts rather than in a sustained fashion (Rusiniak et al., 2018), we propose that alpha-gated re-evaluation mostly occurs within these bursts. That is, in BR the re-evaluation of competing stimuli might not unfold homogenously over time, but intermittently. Incidentally, this may well be the origin of the stochastic dynamics that characterises the BR phenomenon. In our protocol, entrainment at the natural alpha rhythm must have increased alpha burst episodes, explaining the faster alternation rate under IAF, compared to no entrainment. During these alpha bursts, in order to reduce stimulus complexity, information is compressed into discrete packages (Vanrullen & Koch, 2003). The dynamics of the system alternate between intervals where the probability of switch is low and recurring intervals of alpha activity where competition between conflicting representations is steep. Therefore, by increasing alpha burst episodes (their amount and/or duration), the probability of stimulus re-evaluation increases and, as a result, faster alternation rates can be expected. The role of alpha in the gating by inhibition hypothesis (Jensen & Mazaheri, 2010) is compatible with the hypothesis laid out here. According to the former, sensory information is coded in gamma activity embedded within the high excitability windows of slower alpha cycles. One could argue that, in fact, slower alpha cycles may allow for more nested gamma cycles, hence lead to more accumulation of evidence per alpha cycle, predicting a result contrary to what we observed. However, variations in the speed of gamma may compensate for the different number of cycles per second across alpha speeds. Indeed, several findings suggest that the length of gamma cycles vary between subjects, and subjects with faster alpha also present faster gamma (Baumgarten et al., 2018; Muthukumaraswamy et al., 2009). We speculate that the number of gamma bursts per alpha cycle is approximately constant, therefore faster alpha rhythms would allow for an increased number of gamma bursts per second, in line with our hypothesis and compatible with gating by inhibition.

Regarding frequency specificity, Experiment 4 allowed to confirm that the effect is maximal in the alpha range. Although we observed that alternation rate did not return to baseline for the faster frequency, this was expected. High frequency stimulation may not be an effective test for specificity, given that alpha activity could still be entrained as a sub-harmonic of higher frequencies, as some studies suggest (C. S. Herrmann, 2001; Christoph S. Herrmann et al., 2016). The maximum NAR observed in the alpha range is in good agreement with evidence from O’Shea and Crassini (O’shea & Crassini, 1984) thus supporting the existence of a higher bound.

It is possible that the physical stimulation based on contrast changes used here resulted in a modulation of attention, which has a well-known effect on BR (Jan W. Brascamp & Blake, 2012; Zhang et al., 2011). Compatible with our hypothesis, attention has been proposed to periodically sample rivalling stimuli during BR (Davidson et al., 2018), and it is possible that the increase in NAR observed in the arrhythmic stimulation condition is due to a global increase in arousal. Our findings can be framed in terms of existing theoretical models of BR which incorporate attention (Li et al., 2017). According to these models, attention recurrently (or rhythmically, in our hypothesis) regulates the imbalance between the two rival stimuli with a slow time constant, triggered by a mutual inhibition process that has a fast(er) time constant. We can interpret this model in terms of a slow rhythm (alpha) and a fast rhythm. This faster rhythm could be gamma, which has also been found to correlate with BR dynamics in previous studies (Fesi & Mendola, 2015). The concept of alpha discrete sampling can also be framed within Bayesian models of BR (Gershman et al., 2012; Hohwy et al., 2008). It has been proposed that the continuous input of inconsistent stimuli creates a situation in which neither of the stimuli representations has both high prior and high likelihood. As a result, periodic sampling determines which of the two representations sways the perceptual outcome at any given time (Hohwy et al., 2008).

The present findings provide a framework for the interpretation of inter-individual differences in BR dynamics: in this framework, endogenous alpha cycles are a representative time constant for each subject that regulates the competition process between neural populations representative of each stimuli. Alpha acts as a pacemaker for information accumulation mediating the transition to awareness by setting the tempo for the competition process. According to this interpretation, our results are in line with recent studies suggesting that alpha activity may not regulate objective aspects of perception, such as perceptual sensitivity, but awareness and confidence about perceptual properties (Benwell et al., 2017; Lange et al., 2013; Samaha et al., 2017).

## Abbreviations

BR: binocular rivalry
NAR: natural alternation rate
EEG: electroencephalography
IAF: individual alpha frequency
SSVEP: steady-state visual evoked potentials
XCOH: cross-coherence
TOST: two one sided tests

## Data availability

The datasets generated during and/or analysed during the current study are available from the corresponding author upon reasonable request.

## Acknowledgments

This research was supported by the *Ministerio de Ciencia e Innovación* (PID2019-108531GB-I00 AEI/FEDER), AGAUR Generalitat de Catalunya (2017 SGR 1545). This project has been co-funded with 50% by the European Regional Development Fund under the framework of the FEDER Operative Programme for Catalunya 2014-2020, with a grant of 1,527,637.88€. We want to thank Dr. Márta Szabina Pápai and Indre Pileckyte for helping in the piloting of the Experiments and in EEG data collection.

