## Supplementary information for "Alpha fluctuations regulate the accrual of visual information to awareness"

### Supplementary figures

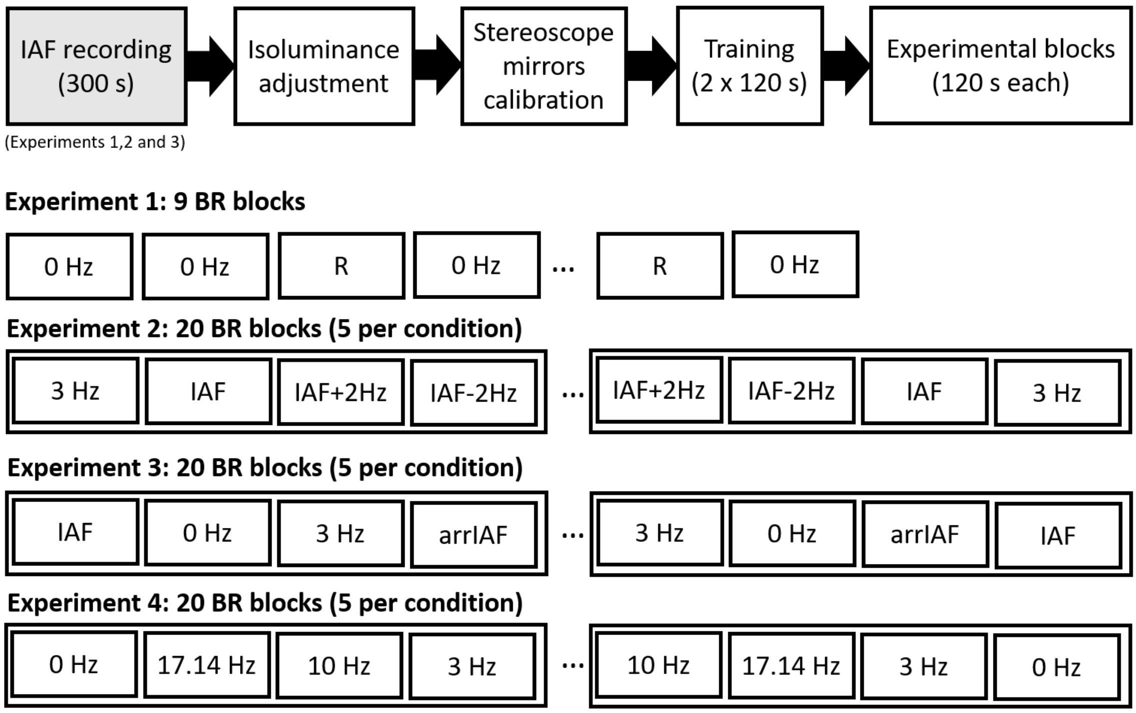


**Supplementary Figure S1: Experimental procedure. All sessions including an electrophysiological recording (Experiments 1, 2 and 3) started with a 5 minutes recording during rest (eyes closed) used to estimate IAF at rest. Next in all the experiments, the RGB level of the green stimulus was adjusted individually and the stereoscope mirrors were calibrated. Prior to the experiment, two training blocks of 2-minute duration were run (0 Hz for experiment 1, random conditions for experiment 2, 3 and 4). Finally, participants ran the experimental blocks. In experiment 1, participants ran 2 blocks of 0 Hz condition and followed by 1 block of Replay (a condition neither used nor relevant for the current experiment) until 9 0 Hz blocks were completed. In experiments 2, 3 and 4 participants performed 5 runs including a block of each of the 4 experimental conditions. The order was assigned randomly, with the constraint that the same condition could not be presented in consecutive blocks. The conditions run in experiment 2 were: 3 Hz, IAF-2Hz, IAF and IAF+2Hz. The conditions run in experiment 3 were: 0 Hz, 3 Hz, IAF and arrIAF. The conditions run in experiment 4 were: 0 Hz, 3 Hz, 10 Hz, 17.14 Hz.**

### Supplementary materials and methods

#### *Sample size calculation*

In Experiment 1, we used data from a previous experiment (1) to estimate the sample size by means of a Monte Carlo randomisation procedure. We sampled with repetition data for sample sizes from 5 to 30 subjects (1000 repetitions). We approximated the statistical power as the proportion of significant correlations observed for a given sample size. The minimum required sample size to obtain a significant true effect 95% of the time for the effect size in (1) was 25 subjects.

In Experiment 2, we used data from Experiment 1 to estimate the effect size of the condition of interest (IAF-2Hz vs IAF+2Hz). First, we estimated that the NAR changed 0.0195 Hz per each Hz in IAF. Consequently, we expected a difference of $\Delta NAR = 0.078$ Hz between slow and fast conditions in Experiment 2. We also assumed that the standard deviation of both slow and fast NAR in Experiment 2 would be equal to the standard deviation in NAR measured in Experiment 1 ($\sigma_{NAR} = 0.069$ ). Then, we used G*Power (RRID: SCR_013726) to calculate the effect size of the difference between matched pairs, assuming uncorrelated populations ($\rho = 0$, which would result in smaller effect sizes).

$$d = \frac{\Delta NAR}{\sigma}, \text{ where } \sigma = \sigma_{NAR}\sqrt{2 \cdot (1-\rho)} \qquad \text{(Eq. 1)}$$

The estimated effect size was d=0.799. Given this effect size d=0.799, the statistical power for a one tailed t-test of two means (matched pairs) is above 95% for 20 subjects, but we selected a sample size of 25 to be able to capture smaller effect sizes (down to d=0.68).

In Experiment 3, there is no previous data comparing rhythmic (IAF) vs. arrhythmic entrainers. In order to have an intuition about the effect size we could expect, we based our calculation on the empirical effect sizes observed in Experiment 2 (N=25), that ranged between d=0.39 (fast vs. slow alpha entrainment) and d=1.88 for the contrast between 3 Hz and IAF+2Hz. We used G*Power (RRID: SCR_013726) to calculate the sample size required to observe a medium effect (d>0.5) with 95% power. We selected 30 participants, which provided 95% statistical power for effects d≥0.62.

In Experiment 4, using the NAR measured for all entraining frequencies (3 to 14 Hz), we fitted a line and we estimated that the change in NAR was 0.02 Hz per each 1 Hz change in entrainment. We estimated that the standard deviation would be the same as the one measured in the previous experiment and we used Eq. 1 to estimate the expected effect size, assuming uncorrelated populations. The estimated effect size was

0.63. The sample size required to measure an effect of 0.63 with 95% statistical power is 29, however, we increased the sample size to 35 to be able to capture effects sizes down to 0.53.

*Inclusion criteria*

NAR estimation stability: we used data from experiment 1 to calculate the CI for the deviation in the estimation of the median with different numbers of percepts, taking as ground truth the value obtained with all the percepts (231±71). For each participant, we performed 1000 random samplings of percepts (without replacement) in order to calculate the average deviation in median estimation. For 50 percepts or more, the deviation in the median estimation decreased to values below 3.5%. We decided to exclude from experiment 2 and 3 subjects with fewer than 50 percepts in at least one of the experimental conditions.

Artefacts in EEG recordings: we controlled the percentage of data discarded by blinks, eye movements or movements in general. In experiment 1, participants with more than 5% of discarded data overall were excluded from the final sample. In experiments 2 and 3, the criterion was raised to 10% discarded data, but applied separately in each of the 4 experimental conditions: participants with more than 10% of discarded data in at least one of the experimental conditions were excluded from the final sample.

*Equivalence interval evaluation*

In order to test the hypothesis that dynamics without entrainment (0 Hz) were equivalent to the dynamics with non-alpha entrainment (3 Hz), we used a Two One Sided Test (TOST)(2), a test specifically designed to reject the presence of effects large enough to be considered worthwhile. This approach separately evaluates two null hypotheses, that if the difference between two conditions falls between a lower and upper bound [$\Delta_L$ $\Delta_U$], it is considered to be equivalent to the absence of an effect. A TOST is considered to be satisfied when the p-value of both left and right-sided tests is below the significant selected level and, in addition, the difference between the two conditions falls within the pre-selected confidence interval.

Therefore, the application of a TOST requires that the lower and upper bounds are selected prior to performing the statistical test. For defining the equivalence interval in experiment 3, we first imposed a symmetric equivalence interval $[\Delta_L\ \Delta_U] = [-\Delta\ \Delta]$ and used data from experiment 1 to determine a reasonable confidence interval. We consider that the difference in two conditions may be attributed to sampling error, so we estimated the magnitude of the differences between NAR from a single population due to sampling error.

We used data from experiment 1 (27 subjects, 9 blocks per subject) to randomly generate 2 datasets (A and B) per subject (4 blocks each). In order to reduce variability between participants, the NAR of each participant's datasets A and B was normalized relative to the NAR of dataset A. We then performed a permutation test, in which we sampled 30 participants from the sample (the sample size selected for experiment 3) with repetition. We performed 1000 random samplings and calculated the proportion of successful TOST at different equivalence intervals: 0 to 20%, in steps of 1%. We selected a 10% equivalence interval, as the statistical power for a sample size of 30 subjects, was above 95%.

*Objective isoluminance adjustment*

We adjusted the RGB value of the green stimulus to subjectively match the luminance of the green stimulus. Previous to the stereoscope mirrors setting and adjustment, a central Gabor was presented alternating between red and green at 60 Hz, and participants adjusted (using up/down keys) the luminance of the green stimulus until the flickering between red and green was minimal (or had disappeared). The procedure was performed twice for each participant, once starting at maximum green luminance and the other starting at minimum green luminance. The mean value between the two was chosen as luminance for the green stimulus for the experiment.

*Visual stimulation in the rhythmic and arrhythmic condition*

In experiment 3, the number of contrast increases in the arrhythmic and rhythmic conditions was very similar: 1265±101 for IAF condition, and 1267±102 for arrIAF condition. Individually, the difference in number of contrast increases ranged from -8 to 6.

We calculated the power spectrum of the visual stimulation for the IAF and arrIAF conditions, using Welch method (5 seconds windows, 10% overlap) at frequencies normalized with respect to stimulation at IAF (from 0 to 4·IAF in steps of 0.025·IAF). As can be seen in supplementary Figure S2, the percentage of power at the IAF in the rhythmic condition (IAF) is around 30%, whereas for the arrhythmic condition (arrIAF), it is around 1%.

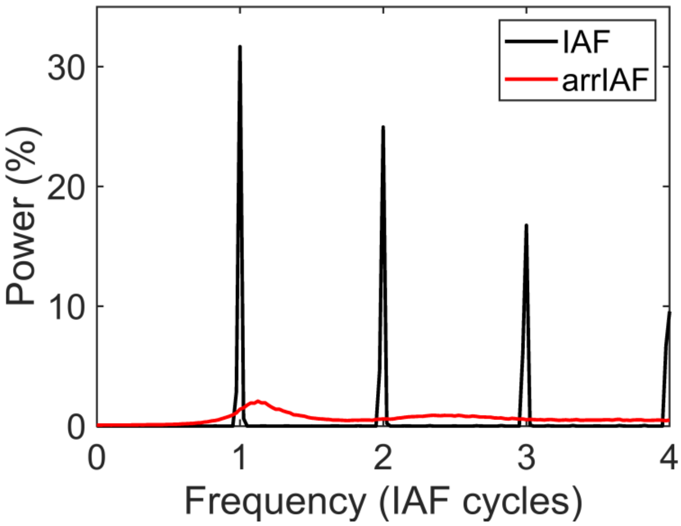


**Supplementary Figure S2: Power spectrum of visual stimulation, expressed as percentage of power in the 0 to 40 Hz band, averaged across participants as a function of normalized stimulation frequency for IAF condition (black) and arrIAF condition (red).**

*Entrainment at single subject level*

At single subject level, the signal to noise ratio of the measurements is smaller than for the average measurements. In order to be able to detect entrained activity at single subject level, we applied Generalized Eigen value Decomposition (GED)(3), a spatial filter that maximized activity in the entrained frequency range.

For each subject and entraining condition (3 Hz, IAF-2Hz, IAF, and IAF+2Hz in experiment 2, 3Hz, and IAF in experiment 3), data from the first 5 seconds preceding the first entrainer was discarded (in order to ensure that the entraining had lasted for several cycles) and the rest was divided into two datasets of equal length (~57.5 s) that, from now on, we will refer as spatial filter dataset and test dataset. For both spatial filter and test datasets, data was averaged across the entraining condition (aligned to the onset of the entrainers).

In order to calculate the spatial filter for each subject and entraining condition, we calculated the signal covariance matrix: time averaged data was demeaned and filtered around the entraining frequency (Gaussian filter, bandwidth 0.5 Hz) and the channel to channel covariance was calculated. This procedure was repeated for building noise covariance matrix, data was demeaned, filtered at frequencies ±1 Hz with respect to entraining frequency (Gaussian filter, bandwidth 2Hz) and channel to channel covariance matrices for both upper and lower filtering were calculated, the average of these two covariance matrices was the noise matrix. Then, we calculated the spatial filter (W): the linear combination of channels that maximized the difference between signal covariance matrix (S) and noise covariance matrix (R) by solving the linear equation:

$$R^{-1}SW = \lambda W$$

Where λ are the eigen values (strength of the eigenvectors) of the eigenvectors in W (each eigen value corresponds to linear combination of channels, the spatial filter). For each subject and entraining frequency, we selected the eigenvector with larger eigen value that presented a parieto-occipital topography and we calculated the power spectrum of the SSVEP for unfiltered test dataset.

**Supplementary results**

*Behavioural characterization*

As expected, the proportion of time spent in each of the percepts was balanced for green and red (Supplementary Figure 3). On average, participants reported a red percept 44±8% of the total recorded time, green 50±8% and no percept 6±8%, with similar proportions for all experimental conditions. For the calculation of NAR, we used an average number of percept per condition 231±71 for experiment 1, 179±66 for experiment 2, 171±85 for experiment 3 and 166±55 for experiment 4. The distributions of percept durations (normalized) for each experimental condition are displayed in Supplementary Figure S3.

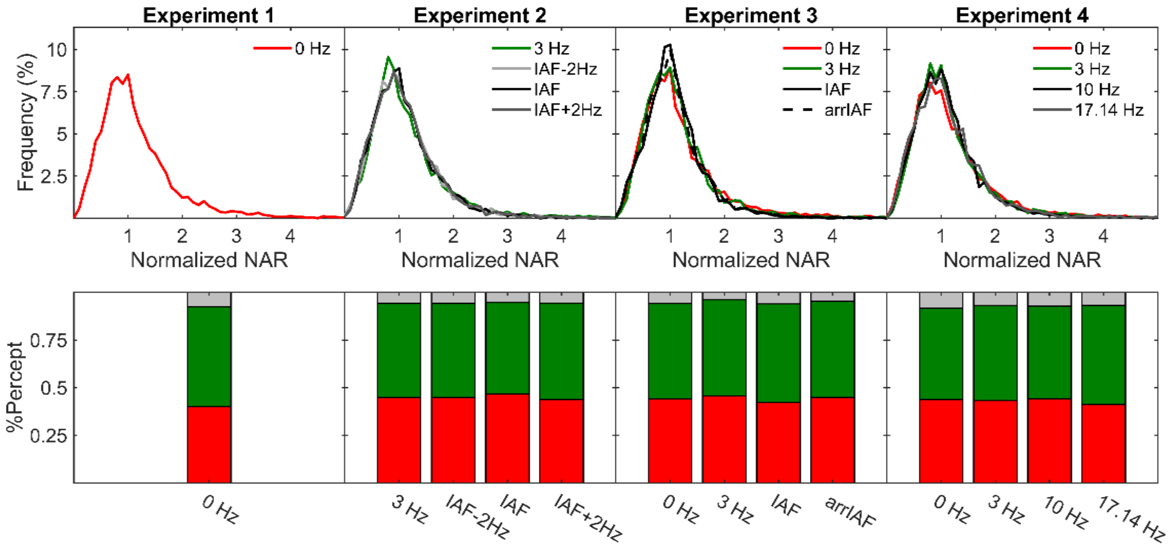


**Supplementary figure S3: Top: normalized NAR collapsed for all participants at experiments (from left to right) 1, 2, 3 and 4. Bottom: average proportion of time spent in each percept (red, green, no percept) for all the experiments and conditions.**

*Reality check: Coefficient of variation*

In Experiments 3 and 4, we performed a post-hoc analysis on the coefficient of variation for each of the entraining conditions. The coefficient of variation (CV) is the ratio between the standard deviation of perceptual alternations and mean alternation time, and it is informative about the interplay between adaptation and noise: values close to 0 indicate that adaptation dominates BR dynamics, whereas values closer to 1 indicate that the noise is driving the competition. We used a TOST (2) to ascertain whether entrainment conditions would result in a different balance between adaptation and competition, compared to no entrainment (0 Hz). The equivalence interval for the CV was determined with a statistical power above 95% for a sample size of 30 subjects. Based on the equivalence interval derived from Experiment 1 (22%), the CV of all experimental conditions was equivalent to the CV of the no entrainment (0 Hz) condition (Supplementary Table 1). This result suggests that the balance between noise and endogenous contributions to BR dynamics was equivalent across conditions.

| **Exp.** | **Condition** | $p_{left}$ | $p_{right}$ | **CI** | **d.o.f.** |
|---|---|---|---|---|---|
| **3** | 3 Hz | $4 \cdot 10^{-5}$ | 0.01 | [-0.05 0.17] | 29 |
| **3** | IAF | 0.03 | 0.003 | [-0.19 0.11] | 29 |
| **3** | arrIAF | 0.009 | 0.0015 | [-0.16 0.10] | 29 |
| **4** | 3 Hz | $10^{-11}$ | 0.002 | [0.03 0.17] | 34 |
| **4** | 10 Hz | $3 \cdot 10^{-6}$ | $7 \cdot 10^{-6}$ | [-0.07 0.08] | 34 |
| **4** | 17.14 Hz | $8 \cdot 10^{-11}$ | $10^{-5}$ | [-0.005 0.12] | 34 |

**Supplementary Table 1: Comparison of the CV for all the experimental conditions in Experiments 3 and 4 with CV at the no entrainment condition (0 Hz). The statistical test performed was a TOST, with an equivalence interval of 22%.**

*Entrainment at single subject level*

The SSVEP at individual level obtained from the GED component with stronger energy

We inspected the SSVEP at individual level, using GED(3) for obtaining spatial filters maximizing entrained activity. For most subjects, we selected the first (stronger) component for calculating SSVEP. In experiment 2, based on the topography of the component, we selected the second stronger component for 3 subjects: 1 subject at 3 Hz, 1 subject at IAF-2Hz and IAF+2Hz and 1 subject at IAF. In experiment 3, for the same reason, we selected the second stronger component for 3 subjects: 1 subject at both 3 Hz and IAF, 1 subject at 3Hz and 1 subject at IAF.

Most subjects (see Supplementary Table 2) displayed a peak at the entrained frequency. This ensures that brain responses were induced at the desired frequencies in the different entrainment conditions, as measured in the occipital electrodes of interest.

| | 3 Hz | IAF-2Hz | IAF | IAF+2Hz |
|---|---|---|---|---|
| **Experiment 2** | 25(0) | 23(2) | 25(0) | 24(1) |
| **Experiment 3** | 30(0) | -- | 27(3) | -- |

**Supplementary Table 2: Number of participants displaying a peak in the frequency of entrainment at individual level in the SSVEP calculated using GED. Between brackets, the number of participants that did not display a peak at the frequency of entrainment in the SSEVP is shown.**

A representative example of the power spectrum of the SSVEP calculated for selected components is shown in Supplementary Figure S4.

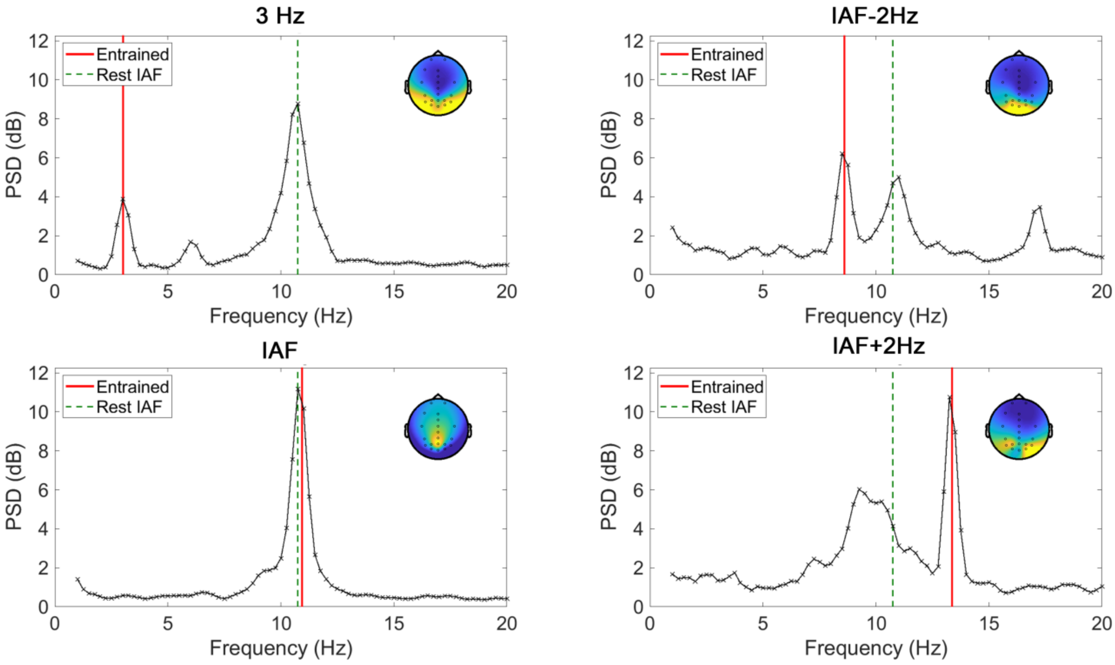


**Supplementary Figure S4: Power spectrum of the SSVEP for a representative subject in experiment 2. For each of the entraining frequencies, the power spectrum displays a clear peak at the entraining frequency (red vertical line); the activity at the individual alpha frequency (green dotted line) is still present. The insets display the topographical distribution of the component selected for SSVEP calculation.**

*Non-planned contrasts*

In Supplementary Table 3 we display the contrasts between conditions that were not planned in our main analysis but may be informative. Notice that we used two tail tests for all non-planned contrasts but for IAF-2Hz vs. IAF and IAF vs. IAF+2Hz, where we used a left tail because we expected a negative difference (slower condition for slower entraining frequency) even if this difference did not reach significance (as experiment was designed for IAF-2Hz vs. IAf+2Hz contrast). All conditions with a high number of visual stimuli result in faster alternation rates than both no visual stimulation and 3Hz stimulation condition.

| Exp. | Cond. 1 | Cond. 2 | Tail | p-value | t-value | d.o.f. | d |
|---|---|---|---|---|---|---|---|
| **2** | IAF-2Hz | IAF | Left | 0.08 | -1.443 | 24 | -0.29 |
| **2** | IAF-2Hz | 3 Hz | Both | $5 \cdot 10^{-9}$ | 7.801 | 24 | 1.56 |
| **2** | IAF | 3 Hz | Both | $7 \cdot 10^{-9}$ | 8.667 | 24 | 1.73 |
| **2** | IAF | IAF+2Hz | Left | 0.12 | -1.223 | 24 | -0.24 |
| **2** | IAF+2Hz | 3 Hz | Both | $1 \cdot 10^{-9}$ | 9.420 | 24 | 1.88 |
| **3** | 0 Hz | IAF | Both | $5 \cdot 10^{-10}$ | 9.108 | 29 | 1.66 |
| **3** | 0 Hz | arrIAF | Both | $6 \cdot 10^{-8}$ | 7.241 | 29 | 1.25 |
| **3** | 3 Hz | IAF | Both | $2 \cdot 10^{-10}$ | 9.420 | 29 | 1.71 |
| **3** | 3 Hz | arrIAF | Both | $5 \cdot 10^{-8}$ | 7.257 | 29 | 1.24 |
| **4** | 0 Hz | 10 Hz | Both | $7 \cdot 10^{-7}$ | 6.037 | 34 | 1.02 |
| **4** | 0 Hz | 17.14 Hz | Both | $9 \cdot 10^{-7}$ | 5.964 | 34 | 1.01 |
| **4** | 3 Hz | 10 Hz | Both | $4 \cdot 10^{-9}$ | 7.835 | 34 | 1.32 |
| **4** | 3 Hz | 17.14 Hz | Both | $1 \cdot 10^{-8}$ | 7.367 | 34 | 1.18 |

**Supplementary Table 3: Statistics for non-planned contrasts in experiments 2, 3 and 4.**

*Phase alignment measures using ITC*

For this analysis, we divided the data in non-overlapping segments of 2 seconds length, locked to a visual stimulus (0.5 seconds to 2.5 seconds after presentation of visual stimulus). Data containing artifacts was rejected, resulting in an average number of trials per condition, 150±31 in experiment 2, and 144±40 in experiment 3. We extracted the phases at the frequencies of interest ( each of the stimulating frequencies) by means of short time Fourier transform (0.5 window length, steps of 2 ms, Hanning window) and calculated inter trial coherence, ITC (4) for each subject condition and electrode in the PO ROI. Next, we averaged the ITC values across the ROI. In order to assess the statistical significance of the phase concentration, we generated surrogate phase distributions for each subject by selecting for each of the available trials, phases at random times in the window of interest. Then, we calculated a surrogate ITC index for each electrode, and repeated the procedure 500 times. Then, we selected for each subject a random surrogate ITC and calculated group level surrogate ITC. This procedure was repeated 10000 times, in order to generate surrogate ITC at group. The p-value of the statistic corresponded to the number of times that the ITC surrogate exceeded the calculated ITC. We used false detection rate to correct for multiple comparisons. After multiple comparison correction, for all the rhythmic conditions we observed significant phase-alignment across the whole window of interest (all p-values<0.0002), whereas for the arrhythmic condition no significant phase concentration was observed (all p-values>0.06).
